# The consensus N_glyco_-X-S/T motif and a previously unknown N_glyco_-N -linked glycosylation are necessary for growth and pathogenicity of *Phytophthora*

**DOI:** 10.1101/2020.05.08.084426

**Authors:** Can Zhang, Shanshan Chen, Fan Zhang, Tongshan Cui, Zhaolin Xue, Weizhen Wang, Borui Zhang, Xili Liu

## Abstract

Asparagine (Asn, N) -linked glycosylation within the glycosylation motif (N_glyco_-X-S/T; X≠P) is a ubiquitously distributed post-translational modification that participates in diverse eukaryotic cellular processes. However, little is known about the characteristic features and roles of N-glycosylation in oomycetes. In this work, it found that 2.5 μg/ml tunicamycin (N-glycosylation inhibitor) completely inhibited *Phytophthora sojae* growth, suggesting that N-glycosylation is necessary for oomycete development. We conducted a glycoproteomic analysis of *P. sojae* to identify and map all N-glycosylated proteins and to quantify differentially expressed glycoproteins associated with mycelia, asexual cysts, and sexual oospores. A total of 355 N-glycosylated proteins were found, containing 496 glycosites that likely participate in glycan degradation, carbon metabolism, glycolysis, or other central metabolic pathways. To verify the glycoproteomic results and further examine the function of N-glycosylation in *P. sojae*, two proteins were selected for PNGase F deglycosylation assays and CRISPR/Cas9-mediated site-directed mutagenesis, including a GPI transamidase protein (GPI16) up-regulated in cysts, with the consensus N_glyco_-X-S/T motif at Asn 94, and a heat shock protein 70 (HSP70) up-regulated in cysts and oospores with a previously unknown N_glyco_-N motif at Asn 270. We demonstrated that the GPI16 and HSP70 are both N-glycosylated proteins, confirming that the N_glyco_-N motif is a target site for asparagine - oligosaccharide N-glycosidic linkage. Glycosite mutations of Asn 94 in the GPI16 led to impaired cyst germination and pathogenicity, while HSP70 mutants exhibited decreased cyst germination and oospore production. This work describes an integrated map of oomycete N-glycoproteomes and advances our understanding of N-glycosylation in oomycetes. Moreover, we confirm that the consensus N_glyco_-X-S/T and the N_glyco_-N -linked glycosites are both essential for the growth of *Phytophthora sojae*, indicating that there are multiple N-glycosylation motifs in oomycetes.

## Introduction

Asparagine (Asn, N) -linked glycosylation is among the most abundant post-translational modifications (PTMs) in a given genome, and is widely distributed across the tree of life (1, 2). This type of modification participates in the regulation of diverse biological processes and reportedly plays a major role in the development of complex multicellular organisms (3, 4). In humans, aberrant glycosylation has been implicated in a number of diseases, including neurodegenerative disorders and cancer (2, 5). Overall, more than 50% of proteins are estimated to be modified by covalent attachment of sugar molecules at some point in their life cycle, higher even than the 30% proportion of proteins predicted to undergo phosphorylation in organisms (6). Due to the complexity of this modification, relatively few large-scale studies of glycosylation have been conducted to date, though development in scope and throughput has increased in recent years. These studies have primarily focused on mammals and lower model organisms such as *Arabidopsis thaliana* and *Saccharomyces cerevisiae* (5, 7, 8). Thus, in order to determine the contributions of N-glycosylation to reproductive fitness and pathogenicity among microorganisms, it is necessary to map N-glycoproteomes in more organisms as well as in more developmental stages of those organisms.

N-glycosylation in the endoplasmic reticulum (ER) is a conserved, two-phase process in eukaryotic cells. First, the assembly of the lipid-linked oligosaccharide (LLO) initiates at the cytoplasmic side of the ER membrane and terminates in the lumen (9, 10). Then, the oligosaccharyltransferase (OST) catalyzes the *en bloc* transfer of the LLO to the asparagine side chain of the acceptor polypeptides in the ER lumen (9, 10). The OST selects polypeptides carrying an N_glyco_-X-S/T (X≠P) motif and generates the N-glycosidic linkage between the side chain amide of asparagine and the LLO (9-12). Moreover, other conserved N-glycosylation motifs have been identified through liquid chromatography - tandem mass spectrometry (LC–MS/MS). For example, the N_glyco_-X-C motif in mice (8), N_glyco_-G and N_glyco_-X-C motifs in yeast (5), as well as the N_glyco_-A, N_glyco_-N and N_glyco_-D motifs in *F. graminearum* (13). However, the verification of these findings and further functional characterization of these N-glycosylation motifs has yet to be conducted.

N-glycosylation affects protein folding either directly or indirectly (14-16). After the protein is correctly folded, three glucose residues are removed and the glycoprotein is exported to the Golgi apparatus for further glycan processing (16, 17). If the N-glycosylation of a polypeptide is disrupted, the protein cannot properly fold into its native conformation, thus indicating the necessity for N-glycosylation in maintaining protein functionality during growth and reproductive development in different mammals and plants species (18-20). Two distinct processes have been described that help eukaryotic cells cope with this problem: ER-associated degradation (ERAD) and the unfolded protein response (UPR) (21). The ERAD system eliminates misfolded proteins via degradation in the cytosol; and the transcriptional induction of UPR target genes allows the cell to adjust the capacity of protein folding in ER (21-23).

Oomycetes classified in the kingdom Stramenopila, are eukaryotic fungal-like microbes (24), many of which, especially *Phytophthora* spp., are important plant pathogens that cause tens of billions of dollars in annual losses to agriculture and aquaculture, globally (25, 26). *Phytophthora sojae* is a widely distributed, destructive oomycete plant pathogen that has been developed as a model for oomycete biology (25, 27). Like most *Phytophthora* species, *P. sojae* reproduces both sexually and asexually. During a disease cycle, the asexual life cycle (mycelium and cyst) is mainly responsible for rapid propagation and dispersal. In contrast, thick-walled oospores produced during sexual reproduction are essential for the pathogen’s long-term survival by introducing greater diversity through recombination and prevention of genetic bottleneck within populations (28-30). Previous research, such as an iTRAQ-based proteomic comparison of mycelia and germinating cysts in *P. capsici* identified several candidate proteins involved in fatty acid degradation for energy production, thereby distinguishing the pre-infection stage from the mycelial stage (31). Proteomic analyses of mature oospores and non-mating hyphae identified proteins enriched in oospores were β-glucanases potentially involved in digesting the oospore wall during germination (32).

Given the relevance of the asexual and sexual phases in the epidemiology of *P. sojae*, we conducted a comprehensive glycoproteomic analysis of the N-glycosylated proteins associated with somatic growth, asexual cysts, and sexual oospores, in order to identify and map the full range of N-glycosylated proteins in this oomycete. In addition, glycoproteomic analysis allowed us to quantify differentially expressed glycoproteins during individual reproductive stages in *P. sojae*. Interestingly, we identified four previously unknown, conserved N-glycosylation motifs in addition to the consensus N_glyco_-X-S/T motif. Furthermore, this work confirmed the N-glycosylation of two highly differentially expressed glycoproteins, including a GPI transamidase protein (GPI16) carrying the canonical consensus N_glyco_-X-S/T target sequence and a heat shock protein 70 (HSP70) carrying a novel N_glyco_-N motif, both of which were shown to be critical for full reproductive capability, and hence pathogenicity in *P. sojae*. To our knowledge, this is the first study to systematically characterize glycoproteins in oomycetes, and to report on function of the N_glyco_-N glycosylation motif.

## Experimental Procedures

### Effect of N-glycosylation inhibitor tunicamycin on *P. sojae*

Tunicamycin is a potent inhibitor of protein N-glycosylation that has frequently been used for experimental chemical inhibition of protein glycosylation in cells (33). Cultures of *P. sojae* strain PS6 were cultivated as described previously (34). The effect of 2.5 μg/ml and 5 μg/ml tunicamycin on *P. sojae* mycelial growth and cyst germination was evaluated using protocols from a previously published study (35). The colony diameters were measured perpendicularly after 6 days dark-incubation, and the germination rate of the cysts was assessed under a light microscope. Tunicamycin was added to the 10% V8 liquid medium following 4 days of incubation of *P. sojae* in the dark at 25°C. Oospore production was quantified 5 days later.

### Preparation of *P. sojae* mycelia, cysts, and oospores

Mycelia were cultured in 100 ml of 10% V8 liquid medium for 4 days at 25°C. Mycelia were then washed with autoclaved, distilled water, then extracted by filtration. Asexual zoospores were obtained following established methods (36). One minute of vigorous shaking of *P. sojae* cultures was used to induce the zoospores to encyst. After harvesting, mycelia and cysts were stored at −80 °C until further use.

Since *P. sojae* is a homothallic species, oospores can be collected from individual colonies after two weeks of dark-incubation on V8 agar at 25°C. The separation of oospores was carried out following the techniques described by Niu (29), with some modifications. Briefly, a Polytron (speed setting 7, Brinkman Instruments, Westbury, NY) was used to homogenize the mating tissues collected from five dishes into 50 ml of sterile deionized water, using five intervals of 2 min each at 4°C. Suspensions of homogenized tissue were then centrifuged at 650 g for 5 min and the supernatant was removed. After filtration through a 100 μm nylon mesh, the hyphae were digested for 1 h in a 20 ml lyase cocktail (0.3 g lyase and 0.12 g cellulase), then washed three times with sucrose solution and deionized water. Finally, the oospore suspension was centrifuged at 4000 X g for 10 min to concentrate the oospores. Oospores were then flash frozen in liquid nitrogen and stored at −80°C.

### Protein extraction and trypsin digestion

The mycelia, cysts, and oospores were thoroughly ground to a fine powder in liquid nitrogen. The powder was sonicated three times on ice using a high intensity ultrasonic processor (Scientz, Ningbo, China) in lysis buffer (10 mM dithiothreitol, 1% protease inhibitor cocktail, and 1% dephosphorylase inhibitor). An equal volume of Tris-saturated phenol (pH 8.0) was then added, and the upper phenol phase was precipitated by the addition of at least four volumes of ammonium sulfate-saturated methanol. After centrifugation at 4°C for 10 min, the remaining precipitate was washed once with ice-cold methanol, followed by three washes in ice-cold acetone. The protein was redissolved in 8 M urea and the protein concentration was determined using a BCA kit (Beyotime Institute, Shanghai, China) according to the manufacturer’s instructions.

Prior to digestion, the protein solution was reduced with 5 mM dithiothreitol for 30 min at 56 °C, then alkylated with 11 mM iodoacetamide for 15 min at room temperature in darkness. The protein sample was diluted by adding 100 mM triethylammonium bicarbonate (TEAB) to adjust the urea concentration to lower than 2M. After this preparation, trypsin was added at a 1:50 trypsin-to-protein mass ratio for the first overnight digestion and at a 1:100 trypsin-to-protein mass ratio for a second 4 h digestion.

### Tandem mass tag (TMT) labeling, high-pressure liquid chromatography (HPLC) fractionation, and N-glycopeptide enrichment

After trypsin digestion, the peptides were desalted using a Strata X C18 SPE column (Phenomenex Torrance, USA) and vacuum-dried. Peptides were reconstituted in 0.5 M TEAB and processed following the instruction manual for the TMT kit (Thermo Fisher Scientific, Waltham, USA). The tryptic peptides were fractionated by high pH reverse-phase HPLC using a Thermo Betasil C18 column (5 μm particles, 10 mm ID, 250 mm length). The hydrophilic interaction liquid chromatography (HILIC) enrichment of N-glycopeptides were performed according to previous reports with minor modifications (37, 38). The peptide fractions were first dissolved in the enrichment loading buffer (80% acetonitrile (ACN); 1% trifluoroacetic acid (TFA)) and then pipetted into a HILIC tip (Jingjie PTM BioLab, Hangzhou, China). After centrifugation at 4000 X g for 15 min, the HILIC tip was washed with 40 μl of loading buffer three times. The enriched N-glycopeptides were eluted with 10% ACN and lyophilized until dry. For deglycosylation, 200 units of PNGase F (NEB) was added into 50 μl of 50 mM NH_4_HCO_3_ in H_2_ ^18^O (Aladdin, Shanghai, Beijing) and incubated at 37°C overnight.

### Liquid chromatography separation and mass spectrometric analysis

Digested peptides were separated using an Acclaim PepMap RSLC C18 capillary reversed-phase analytical column (Thermo) with a 60 min 4-85% ACN/water gradient containing 0.1% formic acid (FA) at a constant flow rate of 700 nL/min with an EASY-nLC 1000 UPLC system (Thermo). The eluted peptides were further ionized and sprayed into the nanospray-ionization (NSI) source followed by tandem mass spectrometry (MS/MS) with a Q ExactiveTM Plus instrument (Thermo) coupled online to UPLC. The electrospray voltage applied was 2.0 kV. The m/z scan range was 350 to 1800 for full scan, and intact peptides were detected in the Orbitrap at a resolution of 70,000. Peptides were then selected for MS/MS using an NCE setting of 28 and the fragments were detected in the Orbitrap at a resolution of 17,500. Automatic gain control (AGC) was set at 5E4.

### Database processing and bioinformatic analysis

The fragmentation data were analyzed using the MaxQuant software package (version 1.5.2.8, http://www.maxquant.org/) (39) to match ions against the *P. sojae* P6497 reference database containing 25721 sequences (downloaded from the Uniprot database, https://www.uniprot.org/). Trypsin/P was specified as the cleavage enzyme, allowing up to 2 missing cleavages. Mass error was set to 5 ppm for precursor ions and 0.02 Da for fragment ions. Carbamidomethylation on Cys (+57.0215 Da) was specified as a fixed modification, while oxidation on Met (+15.9949 Da), and deamidation (^18^O) on Asn (+2.9882 Da) were specified as variable modifications. The false discovery rate (FDR) thresholds for protein, peptide, and modification sites were specified at 0.01.

Soft MoMo (motif-x algorithm, http://meme-suite.org/tools/momo) analysis was used to model the amino acid sequence composition at specific positions in glycosylated 21-mers (10 amino acids upstream and downstream of the site) (40). Secondary structure analysis was performed using NetSurfP (http://www.cbs.dtu.dk/services/NetSurfP/) with *p* values calculated by Wilcoxon test (41). Gene Ontology (GO) terms for the identified proteins were assessed by querying the UniProt-GO annotation database (http://www.ebi.ac.uk/GOA) (42). The protein domain functional annotations were assigned using InterProScan, as implemented in the InterPro domain database (http://www.ebi.ac.uk/interpro/) (43). Soft WolF PSORT version 2.0 (http://www.genscript.com/psort/wolfpsort.html) was used for protein subcellular localization prediction (44).

### Analysis of differentially expressed modified proteins

The ratios of the TMT reporter ion intensities in MS/MS spectra (m/z) from raw data sets were used to calculate fold changes between samples. Whole-proteome analysis was also performed to identify differentially expressed N-glycosylated proteins between the mycelial, cyst, and oospore growth and reproductive stages of *P. sojae*, as described above (except for the enrichment step). The N-glycosylation data were then normalized to the proteome data. In general, the differentially expressed modified and non-modified proteins were defined as having a fold-change ≥ 1.50 or ≤ 0.67, and a *p* value < 0.05.

### Plasmid construction for overexpression and site-directed mutagenesis

To further validate the N-glycosylation modifications identified by proteomic analysis, a GPI transamidase glycoprotein (GPI16) with a consensus N_glyco_-X-S/T motif that was highly expressed in the cysts, and a heat shock protein 70 (HSP70) with a N_glyco_-N motif highly expressed in oospores were cloned. The sequences of the targeted candidate proteins were obtained by searching the *P. sojae* genome, v3.0 (https://genome.jgi.doe.gov/Physo3/Physo3.home.html). The full-length genes were amplified using EasyPfu DNA Polymerase (TransGen Biotech, China) and confirmed by Sanger sequencing (Primers in Table S1). Substitution mutations were introduced into the genes encoding the two candidate proteins by oligonucleotide-directed mutagenesis (Table S1) (45). The oligonucleotides used to alter the potential N-glycosylation sites were designed to replace the codon for asparagine (Asn, N) with that of alanine (Ala, A). After sequencing, no changes were detected other than the desired nucleotide substitutions in the cDNAs of mutant variants.

For protein overexpression to validate the identified glycoproteins, the target genes with or without glycosite mutation were cloned into the pTOR::FLAG vector using the In-fusion HD Cloning kit (Clontech, Beijing, China) (Table S1). The CRISPR/Cas9 system was used for excision and replacement of the wild-type target proteins with glycosite-mutagenized variants in order to better understand the functions of these N-glycosylation sites in *P. sojae* growth and reproduction. Constructs encoding two single guide RNAs (sgRNAs) (sg214 for GPI16: ACTCGGCGTCGTGTTTCTCG; and sg603 for HSP70: CGTCTGGTAGTCCTCCTCCG) were cloned separately into plasmid PYF515. To avoid the sgRNA-mediated Cas9 cleavage of the homology directed repair donor, the sgRNA binding residues in the donor plasmid were mutated without changing the encoded amino acids, producing the two donor plasmids. Flanking sequences, 1,000 bp upstream and downstream of the mutation site, were amplified using GPI16-F1/R1, GPI16-F2/R2, and HSP70-F1/R1, HSP70-F2/R2 primers (Table S1) and ligated into pBS-SK^_+_^ using the In-fusion HD Cloning kit (Clontech, Beijing, China), respectively, to create the basic donor plasmid (46).

### Transformation of *P. sojae*

The transformation of *P. sojae* was accomplished by following protocols described in Fang and Tyler (47). To acquire the overexpression mutants, the pTOR::FLAG plasmids carrying each glycoprotein were respectively transformed into *P. sojae*. PYF515-sgRNA was co-transformed into *P. sojae* PS6 together with its donor plasmid for the site-directed mutagenesis of the two target glycoproteins. The mutants were initially screened in pea broth medium amended with 30 μg/ml geneticin (Amresco, Solon, United States) and then on V8 agar plates containing 50 μg/ml geneticin. The hyphae of the mutants were then collected for genomic DNA (gDNA) extraction. For verification of the mutants, the amplification products of the full-length target candidate genes were confirmed by sequencing in all mutants.

### The validation of target glycoproteins

Harvested hyphae were ground to a powder using a pestle and mortar, and the total protein was extracted using a Protein Extraction kit (Invent, Beijng, China). Cell debris was removed by centrifugation for 10 min at 13000 x g at 4°C. Anti-FLAG M2 beads (Sigma-Aldrich, Shanghai, China) were added to the lysate, followed by incubation for 5 h at 4°C with gentle shaking. Proteins binding to the beads were eluted into elution buffer (7 M urea, 2 M thiourea, 65 mM dithiothreitol and 200 mM PMSF). PNGase F (NEB, Beijing, China) was used for the deglycosylation analysis with an overnight incubation at 37°C.

Proteins were separated in SDS-PAGE gels and the separated proteins were transferred to polyvinylidene difluoride membranes. The membranes were then blocked using PBSTM buffer (phosphate-buffered saline with 0.1% Tween 20 and 5% non-fat milk) for 30 min at room temperature with 60 rpm shaking. Anti-FLAG antibodies (1:5000; Sigma-Aldrich, Shanghai, China) were then added to the PBSTM. After addition of the antibody, the membranes were incubated at room temperature for 3 h then washed three times (10 min each) with PBST buffer. The membranes were then incubated with an HRP-conjugated goat anti-mouse IgG antibody (CWBIO, Beijing, China) at a dilution of 1:5000 in PBSTM at room temperature for 1 h with 60 rpm shaking, and followed by three washes (5 min each) with PBST. The signals were detected by Azure c600 imager (Azure Biosystems, USA).

### Phenotypic analysis of the site-directed mutants

Mycelial growth of *P. sojae* mutant strains was assessed by measuring the colony diameter in two perpendicular directions after 6 days of incubation in the dark on 10% V8 agar. Sporangia and zoospores were obtained using previously published protocols (48). Zoospore encystment was measured by pipetting 100 μl of zoospore suspension onto a glass plate and counting the number of encysted zoospores after 4 h incubation at 25°C and 80% humidity. Oospores of *P. sojae* were collected after two weeks of incubation in the dark on V8 agar at 25°C. The 4-day-old etiolated soybean seedlings (*cv*. Ribenqing, grown in the dark at 25°C and 80% humidity) were used for the *in planta* pathogenicity assays (36). The seedlings were spot inoculated at 2 cm below the beginning of the root zone with 10 ul zoospores suspension (1×10^4^ zoospores/ml), and incubated at 25°C for 2 days in the dark. The lengths of the disease lesions above and below the point of inoculation on the hypocotyl were then measured. Each treatment contained at least 10 seedlings and the entire experiment was conducted at least twice.

### RNA extraction and Quantitative real time PCR assays

Quantitative real time PCR (qRT-PCR) was used to assess the relative expression levels of the genes in the ER-associated degradation (ERAD) pathway and the unfolded protein response (UPR) pathway in the glycosite mutant strains and wild-type *P. sojae*. The biological samples were prepared according protocols established in the above section. Total RNA was first extracted using the SV Total RNA Isolation kit (Promega, Beijing, China) before cDNA was synthesized using the PrimeScript RT reagent kit with gDNA Eraser (Takara, Beijing, China) following the protocols provided by the manufacturer. The qPCR reactions were performed using the SYBR Premix Dimer Eraser kit (Takara, Beijing, China). The relative quantities of the PCR products were calculated using the 2^-ΔΔCt^ method, with the *RPS* and *RPL13a* genes used as internal references to normalize the transcript levels. The primers used in the qRT-PCR experiments are listed in Table S1. All treatments were represented by at least three replicates and the entire experiment conducted three times.

## Results

### N-glycosylation inhibitor tunicamycin significantly inhibited growth of *P. sojae*

The N-glycosylation inhibitor tunicamycin is used extensively for manipulation or prevention of protein N-glycosylation in cells (33), although its effects on oomycetes have not been reported. We first examined the effect of tunicamycin on mycelial growth, cyst germination, and oospore production in *P. sojae* and found that 2.5 μg/ml tunicamycin completely inhibited the growth of *P. sojae* (Fig. 1A). The rate of cyst germination dropped from 81.5% in the untreated control to 10.8% and 7.6% following the addition of 2.5 μg/ml or 5 μg/ml tunicamycin, respectively (Fig. 1B). The WT *P. sojae* exhibited a slight decrease in oospore production under exposure to 2.5 μg/ml tunicamycin, but a significant decrease under treatment with 5 μg/ml tunicamycin (*P* < 0.001, Fig. 1C). These results indicated that N-glycosylation likely plays an essential role in the growth of *P. sojae*.

**Fig. 1.**
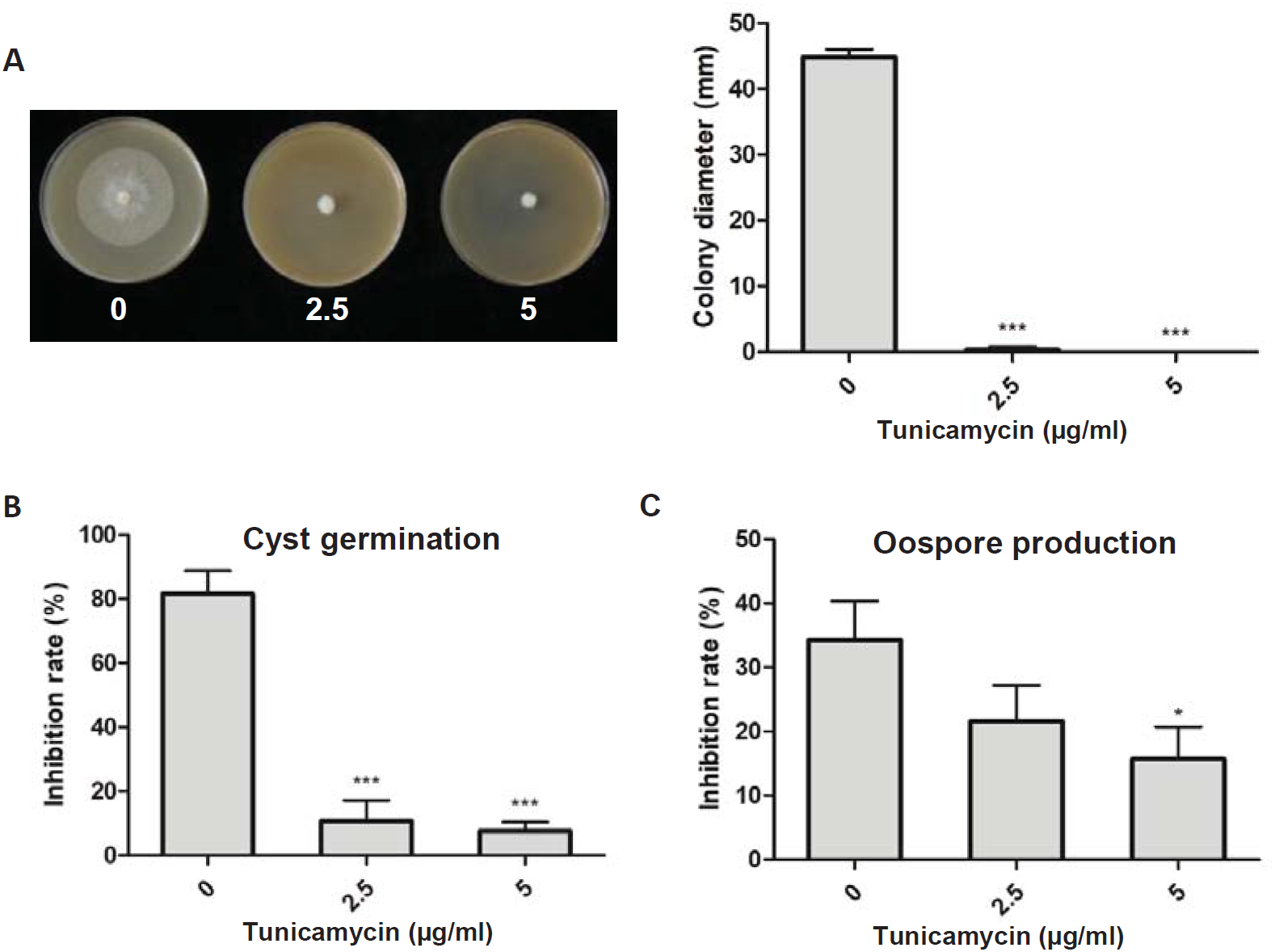
Effect of N-glycosylation inhibitor tunicamycin on the growth of *P. sojae*. 2.5 μg/ml and 5 μg/ml of tunicamycin was added to medium, and changes in *P. sojae* colony size, cyst germination, and oospore production were recorded. (A) The colony diameters of *P. sojae* cultured on V8 medium for five days. (B) Cysts were treated with tunicamycin germination was determined after 8-12 hours. (C) Tunicamycin was added to the 10% V8 liquid medium following 4 days of incubation of *P. sojae* in the dark at 25°C. Oospore production was determined after 5 days of incubation. Bars represent the mean ± standard error (SD) from at least three biological replicates, and three repeated experiments. An asterisk indicates significant differences based on unpaired Student’s *t* test with the *P* values marked (\**P* < 0.05; \*\**P* < 0.01, \*\*\**P* < 0.001).

### A large set of glycosylation sites were identified across a range of proteins in *P. sojae*

In order to obtain a comprehensive view of N-glycosylation patterns in *P. sojae*, we conducted tandem mass spectrometry on the enriched N-glycopeptides from mycelia, cysts, and oospores. A total of 474 N-glycopeptides were identified with high confidence, corresponding to 355 proteins and 496 sites, using Maxquant software to search our MS/MS fragments against the *P. sojae* P6497 reference proteome. The false discovery rate (FDR) for both glycopeptides and glycosylation sites was adjusted to <1%. Among the identified N-glycosylation sites and proteins, we quantified the glycosylation level, or number of modified sites per protein for 341 sites across 239 proteins in the three different developmental stages (Fig. S1A). To our knowledge, this dataset represents the first oomycete quantitative N-glycoproteome to date.

The mass accuracy distribution of the identified glycopeptides (Fig. S1B) showed that all of the peptides had a mass error of < 5.5 ppm, supporting the high confidence of their identification. A majority of the N-glycosylated proteins contained between one and three post-translational modification (PTM) sites. Specifically, 71.27% of the 355 proteins had one N-glycosylation site, whereas 20.00% and 7.32% of the PTM proteins had two and three N-glycosylation sites, respectively (Fig. S1C). The number of modified proteins with four or more PTM sites was significantly lower. The alpha-mannosidase protein (accession No: G4Z5K4) had the highest number (six) of N-glycosylation sites, while the tyrosinase protein (accession No: G5A886) had five sites. We thus found and identified a large number of N-glycosylated proteins, indicating that N-glycosylation is widely present in *Phytophthora*. These glycoproteins showed differences in the number of PTM sites, suggesting that N-glycosylation may not affect each glycoprotein equally.

### N-glycosylation preferentially located on five conserved N-glycosylation motifs

To identify sequence motifs that were generally associated with N-glycosylation sites, we examined a 21-amino acid sequence window surrounding the central N_glyco_ residues of each site using the web-based Motif-X program with a significance threshold of *P* < 0.000001. Previous research showed that the consensus sequence for N-glycosylation motif is N_glyco_-X-S/T (X≠P) (10-13). In this study, we found that most of the 248 N-glycosylation sites matched the N_glyco_-X-S/T motif (Fig. 2A). Interestingly, four other conserved motifs were identified among the regions surrounding N-glycosylation peptides, including N_glyco_-A, N_glyco_-N, N_glyco_-D, and N_glyco_-S (Fig. 2A). As indicated by the heat map of the frequencies of different amino acids that were adjacent to the N_glyco_ residues (Fig. 2B), N-glycosylated sequences were non-randomly distributed among residues, with higher frequencies associated with glycine (G) at the −1, −6 site and lysine (K) at the −7, −8 site. Three nonpolar residues, including proline (P), leucine (L), and phenylalanine (F), more frequently occupied the +1 or +2 site (the cut-off value for the log_10_ transformed Fisher’s exact test *P* value was set at 1).

**Fig. 2.**
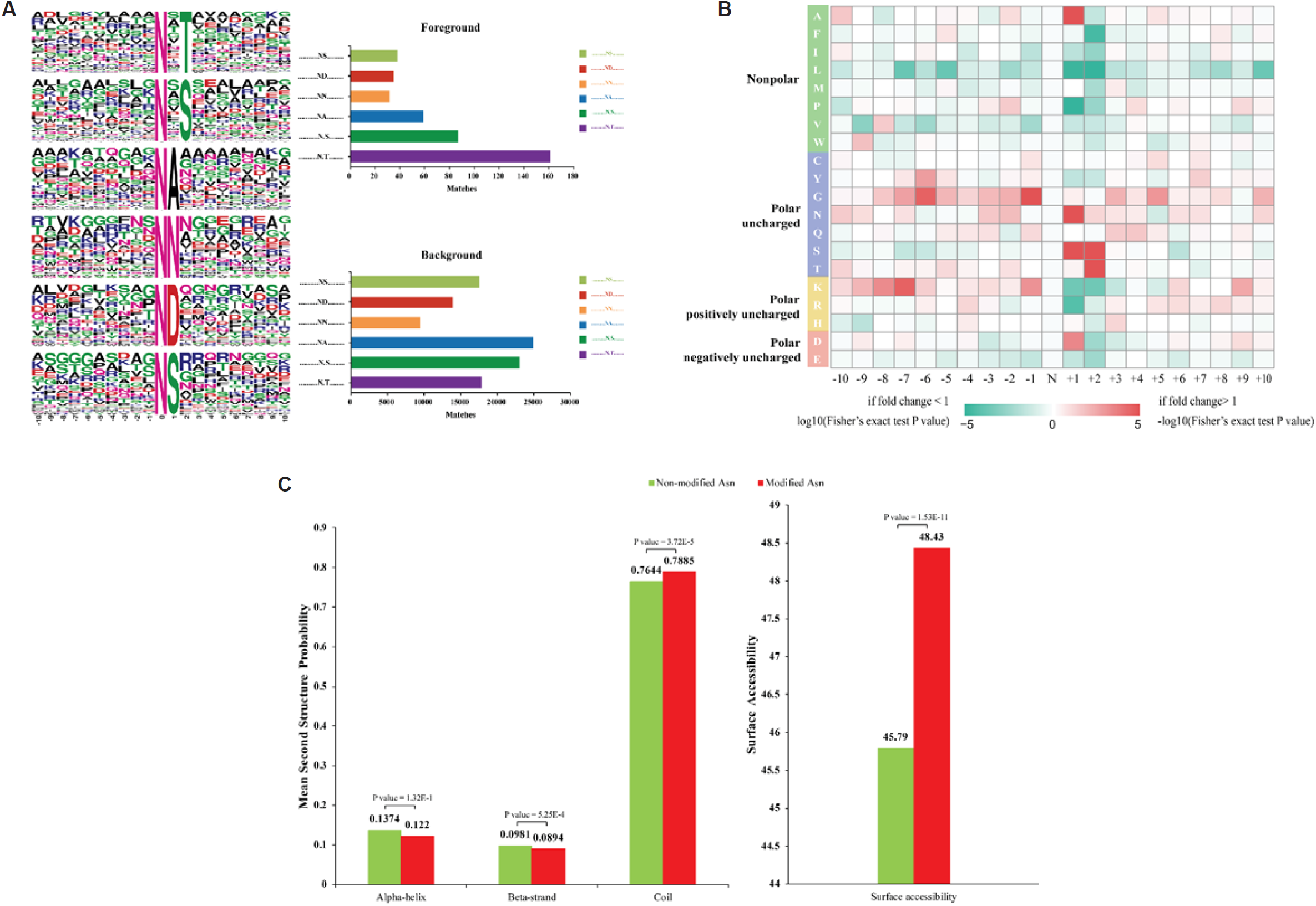
Properties of N-glycosylated peptides in *P. sojae*. (A) Motif analysis of N-glycosylation peptides in *P. sojae*. Sequence models comprising amino acids in specific positions of modified-21-mers (10 amino acids upstream and downstream of the site) of all of the protein sequences. All of the database protein sequences were used as the background database parameter. Both the foreground and background histograms are shown on the left; (B) Heatmap showing the frequency of different amino acid residues around glycosylated Asn residues in *P. sojae*; (C) Distribution of glycosylated Asn and non-glycosylated Asn in protein secondary structures. The mean secondary structure probabilities of modified Asn residues were compared against the mean secondary structure probabilities of the dataset containing non-modified Asn residues from all N-glycosylated proteins identified in this study. *P* values were calculated using non-paired Wilcox tests.

To further understand the effects of N-glycosylation on protein structure, secondary structural predictions were performed using the NetSurfP algorithm on both the modified subset and un-modified Asn residues. We observed that N-glycosylation was more likely to occur on the coil structures (*P* value = 3.72 E-5). In contrast, N-glycosylated residues were less likely to be located on beta-strands (*P* value = 5.25 E-4), and no significant differences were found between modified and un-modified Asn residues on alpha-strands (*P* value = 1.32 E-1) (Fig. 2C, left). The surface accessibility of N-glycosylation sites was significantly higher than that of un-modified Asn residues (*P* value = 1.53 E-11), indicating that N-glycosylated residues may be preferentially located on the outside of the modified proteins (Fig. 2C, right). This result may also explain the predominant co-localization with hydrophilic amino acids near the N_glyco_ motifs.

### *P. sojae* glycoproteins belonged to a wide range of biological processes

To clarify the functions of these N-glycosylation proteins in *P. sojae*, all of the identified N-glycosylated proteins were subjected to subcellular localization prediction, GO functional classification, and KEGG pathway analyses. Predictions of subcellular localization suggested that among the identified glycoproteins in mycelia, cysts, and oospores, 29.58% were targeted for extracellular secretion, 20.28% were localized in mitochondria, 19.72% were distributed in the cytoplasm, and 14.37% were targeted to the plasma membrane (Fig. 3A). Moreover, most of the glycoproteins matched the N_glyco_-X-S/T motif were more likely to be extracellular (Fig. 3B). However, the glycoproteins carrying four other conserved N-glycosylation motifs, including N_glyco_-A, N_glyco_-N, N_glyco_-D, and N_glyco_-S, preferred to localize in mitochondria, with a higher percentage of 28.93% than in other localizations (Fig. 3C).

**Fig. 3.**
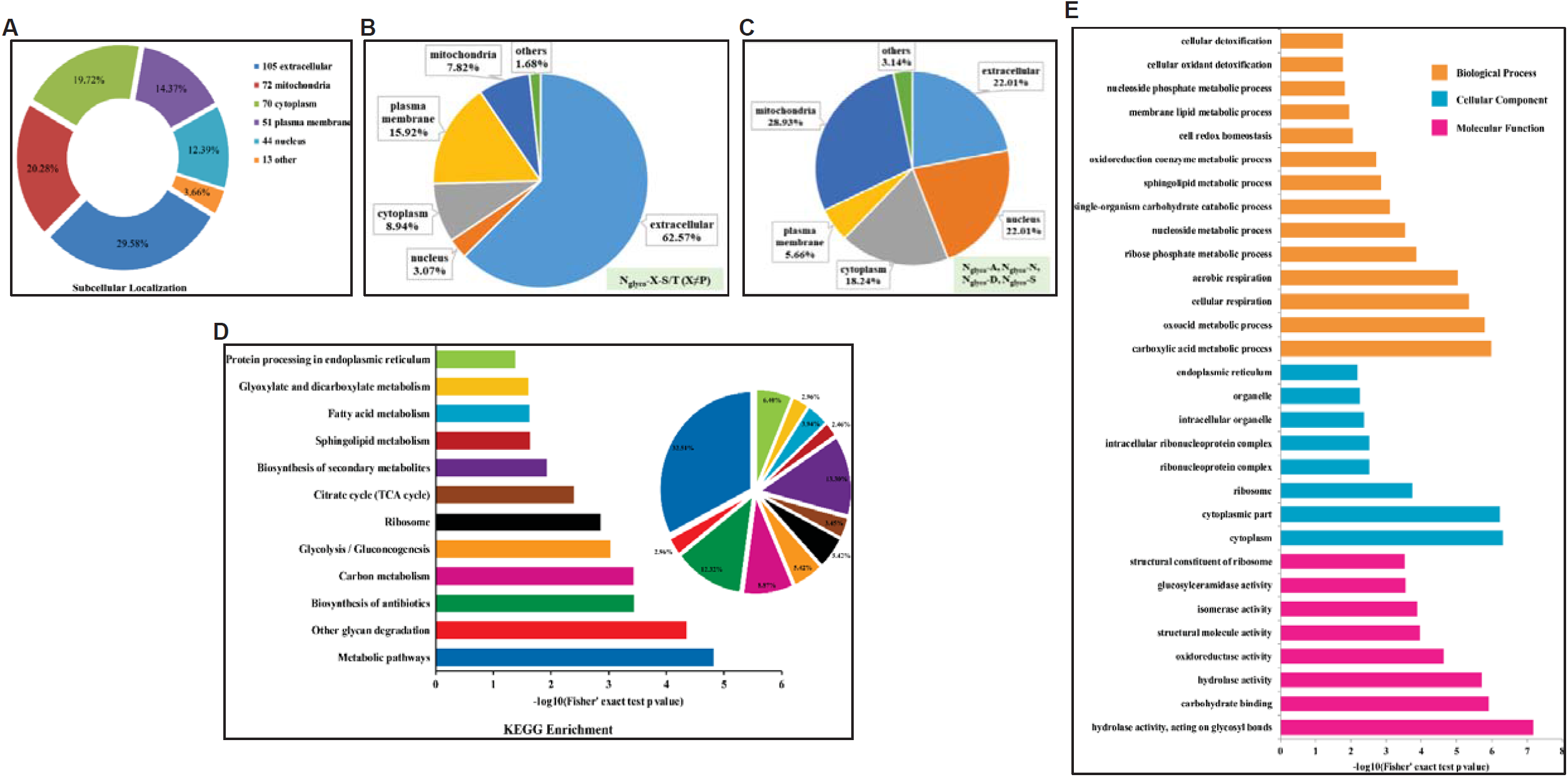
Subcellular prediction and functional classifications of N-glycosylation proteins in *P. sojae*. (A) Subcellular localization of the identified N-glycosylation proteins represented as a pie chart; (B) Subcellular localization of the N-glycosylation proteins carrying N_glyco_-X-S/T motif represented as a pie chart; (C) Subcellular localization of the N-glycosylation proteins carrying N_glyco_-A, N_glyco_-N, N_glyco_-D, and N_glyco_-S motifs represented as a pie chart; (D) GO-based enrichment analysis of identified proteins in *P. sojae*. A two-tailed Fisher’s exact test was employed to test the enrichment of the identified proteins in each category against all of the database proteins. GOs with corrected *P* values < 0.05 were considered significantly enriched; (E) KEGG pathway-based enrichment analysis of N-glycosylated proteins in *P. sojae*. Analysis of the enriched, identified proteins compared to all KEGG database proteins was used to determine enriched pathways via two-tailed Fisher’s exact tests. Pathways with corrected *P* values < 0.05 were considered significantly enriched.

Previous studies have demonstrated that post-translational N-glycosylation is widely distributed throughout many important central metabolic pathways (49). Thus, to better understand the general functions of glycoproteins identified in *P. sojae*, the GO and KEGG database were used to explore the specific metabolic pathways associated with these N-glycosylated proteins. Within the three GO categories (biological process, molecular function, and cellular component), N-glycosylated proteins were assigned to a wide range of biological process sub-categories, including carboxylic acid metabolic process, aerobic respiration, nucleoside metabolic process, single-organism carbohydrate catabolic process, and membrane lipid metabolic process (Fig. 3D). KEGG analysis revealed that the N-glycosylated proteins were involved in glycan degradation (2.96%), biosynthesis of antibiotics (12.32%), carbon metabolism (8.87%), glycolysis/gluconeogenesis (5.42%), ribosome (5.42%), TCA cycle (3.45%), and protein processing in the endoplasmic reticulum (6.40%) (Fig. 3E). These results suggest that *P. sojae* may directly regulate important metabolic pathways through N-glycosylation, especially for carbon metabolism, aerobic respiration, and nucleoside metabolic processes, as well as other functional proteins.

### Changes in N-glycosylation profiles across three developmental stages

We analyzed the differentially expressed N-glycosylated proteins (DEPs) with normalization of the proteome data in the three samples of *P. sojae*. Among the 239 glycoproteins, 89 of them, containing 106 glycosites, showed differences in abundance between cysts and mycelia (Fig. S2A). Moreover, 77 DEPs were common to both oospore comparison groups (Oo vs Cy; Oo vs My), while only 32 or 24 DEPs were shared in cyst comparisons (Cy vs My; Oo vs Cy) or mycelium comparisons (Cy vs My; Oo vs My), respectively (Fig. S2B). These results suggested that in *P. sojae*, the diversity, or the number of different DEPs, was greater during oospore life stages. Based on GO, KEGG, and domain pathway enrichment analysis, N-glycosylated proteins associated with iron binding and protein transport were highly enriched in oospores (Fig. S3). However, a higher proportion of these proteins in the Cy vs My comparison could not be functionally clustered, reflecting the fact that many of the proteins associated with cyst development have yet to be characterized in *P. sojae* and other related organisms.

Two significantly differentially expressed glycoproteins with different N-glycosylation motifs were selected to first validate the glycoproteomic data, and second to examine the contribution of glycosylation to protein function in reproductive development and pathogenicity. These proteins included a GPI transamidase component protein (GPTI, accession No: G4Z0S6) enriched in the cyst stage and carrying a consensus N_glyco_-X-S/T motif, and a heat shock protein 70 (HSP70, accession No: G5A5V6) carrying a novel N_glyco_-N motif that was up-regulated in the cyst and oospore stage (Table 1). The mass spectra used for N-glycosylation identification of these three glycopeptides are shown in Fig. S4. The GPTI glycoprotein only had one glycosite at Asn 94, while the HSP70 protein had two glycosites: one at Asn 270 and the other at Asn 305 (Table 1). The expression level of GPTI-Asn 94 was 2.00-fold higher in cysts than in mycelia and 5.56-fold higher than in oospores, respectively. Similarly, the expression of HSP70-Asn 270 was 1.38- and 1.20-fold higher in cyst and oospores than in mycelia, respectively. Since HSP70-Asn 305 showed no significant up-regulation in either cyst or oospore stage, this Asn residue glycosite was not included in further experiments (Table 1).

**Table 1.** Annotation and recognition sequences of two candidate glycoproteins with increased abundance in cysts and oospores.

| Protein accession<sup>a</sup> | Protein description | Protein Name<sup>b</sup> | Position<sup>c</sup> | Modified sequence<sup>d</sup> | Cy vs My<sup>e</sup> | Oo vs Cy | Oo vs My |
| --- | --- | --- | --- | --- | --- | --- | --- |
| G4Z0S6 | GPI transamidase component | GPI16 | **94** | YGSAQIFN(**1**)R | 2.00 | 0.18 | 0.35 |
| G5A5V6 | Heat shock 70 kDa protein | HSP70 | **270**<br>305 | AETTISN(**0.997**)N(0.003)R<br>N(**1**)DLEQFIYR | 1.38<br>1.13 | 0.87<br>0.57 | 1.20<br>0.65 |

<sup>a</sup> The accession number of candidate proteins in JGI database. <sup>b</sup> The abbreviated name corresponding to each glycoprotein used in this study. <sup>c</sup> Position means the N-glycosylation site of each candidate protein. <sup>d</sup> The number after the amino acid Asn (N) indicates the localization probability of each N-glycosylation site. <sup>e</sup> The comparison ratio of N-glycosylation levels in the two comparison groups.

### Two candidate glycoproteins carrying different conserved glycosylation motif were verified

To verify N-glycosylation of the two candidate glycoproteins, the proteins were checked by immunoblot following treatment with peptide-N-glycosidase F (PNGase F), which catalyzes the removal of N-linked oligosaccharide chains from glycoproteins (50). The results showed that the apparent molecular weight of GPI16 and HSP70 were clearly lower after PNGase F deglycosylation, indicating that both proteins were originally glycosylated (Fig. 4A and Fig. 4C). To further characterize the N-glycosylation sites, GPI16 protein was mutagenized by point mutation at glycosite Asn 94 (GPI16^N94A^) and HSP70 was given a point mutation at glycosite Asn 270 (HSP70^N270A^), then checked again by immunoblotting. As expected, GPI16^N94A^ and HSP70^N270A^ could be separated from GPI16 and HSP70, respectively; and the molecular weight became smaller compared to each parental wild-type protein (Fig. 4B and Fig. 4D). This result demonstrated that the Asn 94 consensus N_glyco_-X-S/T motif and Asn 270 N_glyco_-N motif are both glycosites, and more than one conserved glycosylation motif was found in *P. sojae*.

**Fig. 4.**
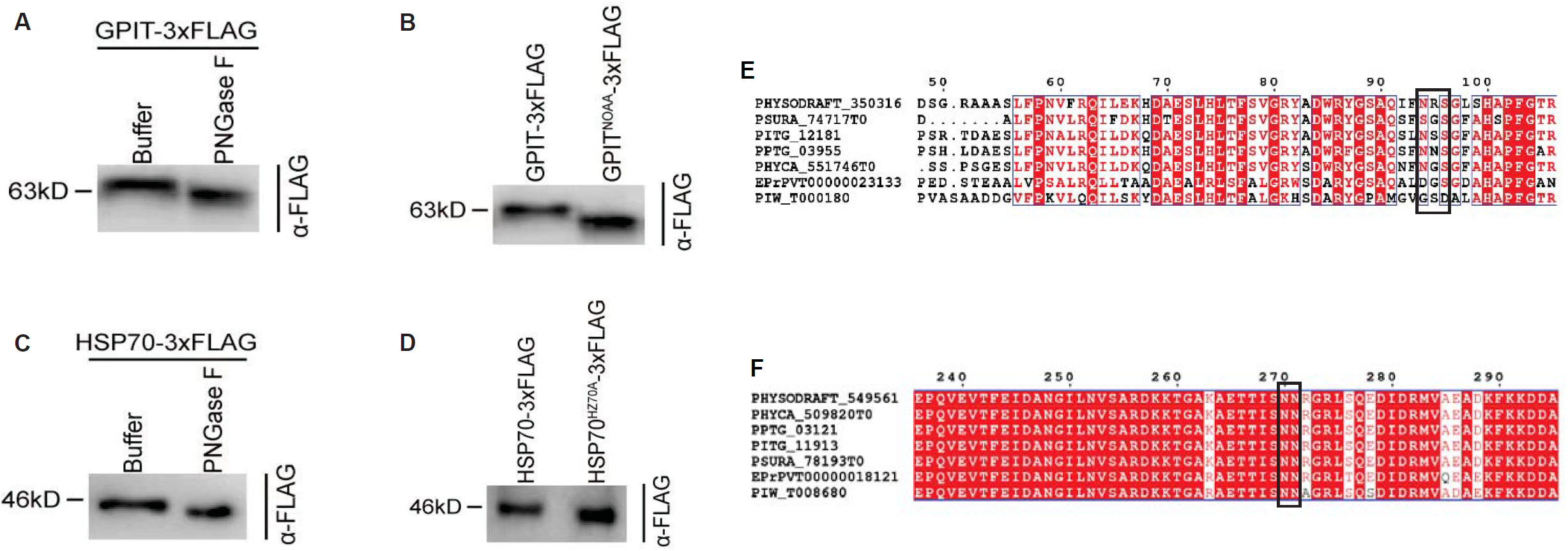
The GPI transamidase (GPI16) with a N_glyco_-X-S/T motif and heat shock protein 70 (HSP70) with a N_glyco_-N motif are both N-glycosylated in *P. sojae*. Total protein extracts were treated with PNGase F or buffer only before immunoblotting; Anti-FLAG antibodies were used for GPI16 (A) and HSP70 (B). Deglycosylation analysis was conducted for the GPI16^N94A^ and HSP70 ^N270A^ mutants, each carrying a point mutation at the Asn 94 for GPI16 (C) and Asn 94 for HSP70 (D) respective glycosylation sites. The conservation of glycosylation site on the two target proteins GPI16 (E) and HSP70 (F) were analyzed in oomycetes, as indicated by PHYSODRAFT for *P. sojae*, PSURA for *P. ramorum*, PTPG for *P. infestans*, PPTG for *P. parasitica*, and PHYCA for *P. capisici*, EPrPVT for *Pythium vexans*, and PIW for *Py. iwayamai*, respectively.

The N-glycosylation motifs of GPI16 and HSP70 were further analysed in different oomycetes species, including four *Phytophthora* species, and two *Pythium* species. The N-glycosylation motifs of these two target proteins, no matter the N_glyco_-X-S/T or the N_glyco_-N motif were both conserved in *Phytophthora* species, as indicate by PSURA for *P. ramorum*, PTPG for *P. infestans*, PPTG for *P. parasitica*, and PHYCA for *P. capisici* (Fig. 4E and Fig. 4F). In addition, only the N_glyco_-N motif of HSP70 was conserved among *Pythium* species, while the motif of GPI16 did not appear in *Pythium* species at the corresponding position (Fig. 4E and Fig. 4F). The conservation of glycosylation sites in oomycetes, especially in *Phytophthora*, suggest they might share similar important functions in oomycetes.

### N-glycosylation of the target glycoproteins is necessary for the growth of *P. sojae*

In order to investigate the function of significantly differentially expressed N-glycosylation site(s) in the two candidate glycoproteins, the CRISPR/Cas9 system was used to replace the Asn residue target site with an Ala residue. We characterized and compared the phenotypes of the wild-type PS6 (WT), a *P. sojae* control transformed with the empty vector (EV), and *P. sojae* isolates containing mutations that prevent Asn target site glycosylation for each of the glycoproteins. No effect was found on mycelial growth and oospore production on GPI16 mutants (Fig. 5A and Fig. 5C), whereas GPI16 mutations significantly affected the germination rate of cysts and pathogenicity of the mutant isolates against soybean (*P* < 0.001, Fig. 5B and Fig. 5D). The cyst germination rate was 82.7% and 78.5% in the WT and EV isolates, respectively, but only 14.8 and 19.2% in GPI16 homozygous mutants M16 and M29, respectively (Fig. 5B). Additionally, etiolated soybean seedlings inoculated with WT and EV zoospores exhibited a lesion length between 66.8-70.0 mm, while the mutants produced significantly shorter lesions of 48.6-49.6 mm in the mutants (Fig. 5D).

**Fig. 5.**
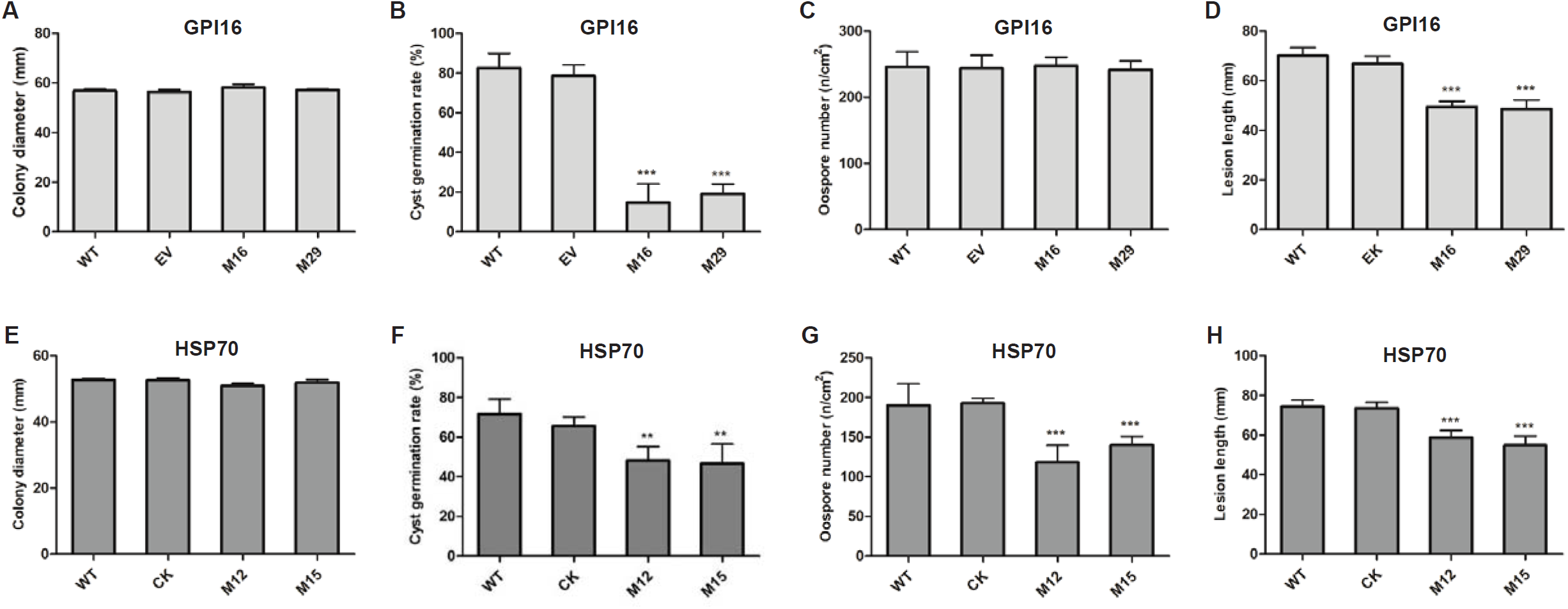
The characteristics of *P. sojae* mutants compared to the wild-type controls. The parental wild-type isolate was PS6; EV was the control isolate of *P. sojae* transformed with empty vector; M16 and M29 were mutants of GPI16 with a point mutation at the Asn 94 (A-D) glycosylation site; M12 and M15 mutant lines carried an HSP70 variant with a point mutation in the Asn 270 glycosylation site (E-H). The mycelial growth (A and E), rate of cyst germination (B and F), the oospore production (C and G), and lesion length on soybean seedlings (D and H) in mutant, WT or control isolates. Bars represent the mean ± standard error (SD) from three biological replicates, and three repeated experiments. An asterisk indicates significant differences based on unpaired Student’s *t* test with the *P* values marked (\**P* < 0.05; \*\**P* < 0.01, \*\*\**P* < 0.001).

The HSP70 mutants showed similar colony diameter as the WT and EV isolates (Fig. 5E). The cyst germination rate was significantly decreased in the HSP70 homozygous mutants compared to WT and EV isolates (*P* < 0.01; Fig. 5F). We observed a significant drop in the average number of oospores from 189.9 to 118.2 and 139.8 in each cm^2^ zone among the mutants M12 and M15, respectively (Fig. 5G). Moreover, pathogenicity was also affected in the HSP70 homozygous mutations (*P* < 0.001, Fig. 5H). In brief, these results of site-directed Asn glycosite mutagenesis were consistent with the relative expression assays indicating low expression of glycosites during developmental stages where mutagenesis had little effect, and high expression in stages that were severely impaired by abolishing glycosylation. This finding further supports the necessity for N-glycosylation of these target proteins for normal growth of *P. sojae*, for both the N_glyco_-X-S/T sequence and the N_glyco_-N motif.

### Elimination of N-glycosylation induces expression of the ERAD and UPR pathway genes

The inhibition of protein N-glycosylation leads to the accumulation of unfolded proteins in the ER (ER stress), and induces expression of the unfolded protein response (UPR) and ER-associated degradation (ERAD) pathway target genes to adapt to the decreased stability of glycoproteins (21-23). We then sought to determine whether UPR and ERAD pathway gene expression was modulated by these mutagenized DEPs. Using qRT-PCR to measure the transcript levels of major ER chaperones and foldases, including *CNE1-2* (accession No: G4YMY1 and G4YVS5), encoding the *P. sojae* homolog of calnexin; *PDI1-3* (accession No: G4YXR9, G4YKG9, G5A3Y0), encoding protein disulfide isomerase; and *ERAD1-5* (accession No: G4Z2L8, G4ZMP2, G4Z0Q7, G4YIZ4, G4ZYS3), encoding ER-associated degradation proteins in WT, EV, and glycosite mutant strains. The expression of two calnexin genes *CNE1-2* was not affected in both GPI16 mutant M16 and HSP70 mutant M12 (Fig. 6A and Fig. 6B). The mRNA levels of *PDI1* and *PDI3* were approximately 1.67- and 1.89-fold higher in the M12 mutant compared with WT, respectively (Fig. 6C and Fig. 6E). In the GPI16 mutant, the expression levels of five *ERAD* genes were all up-regulated, and *ERAD1* appeared 3.03-fold up higher than WT (Fig. 6F-6J). These results indicated that abnormal growth of *P. sojae* caused by elimination of N-glycosylation in these target glycoproteins was potentially related to aberrant protein folding, with different UPR and ERAD genes are playing roles in this process.

**Fig. 6.**
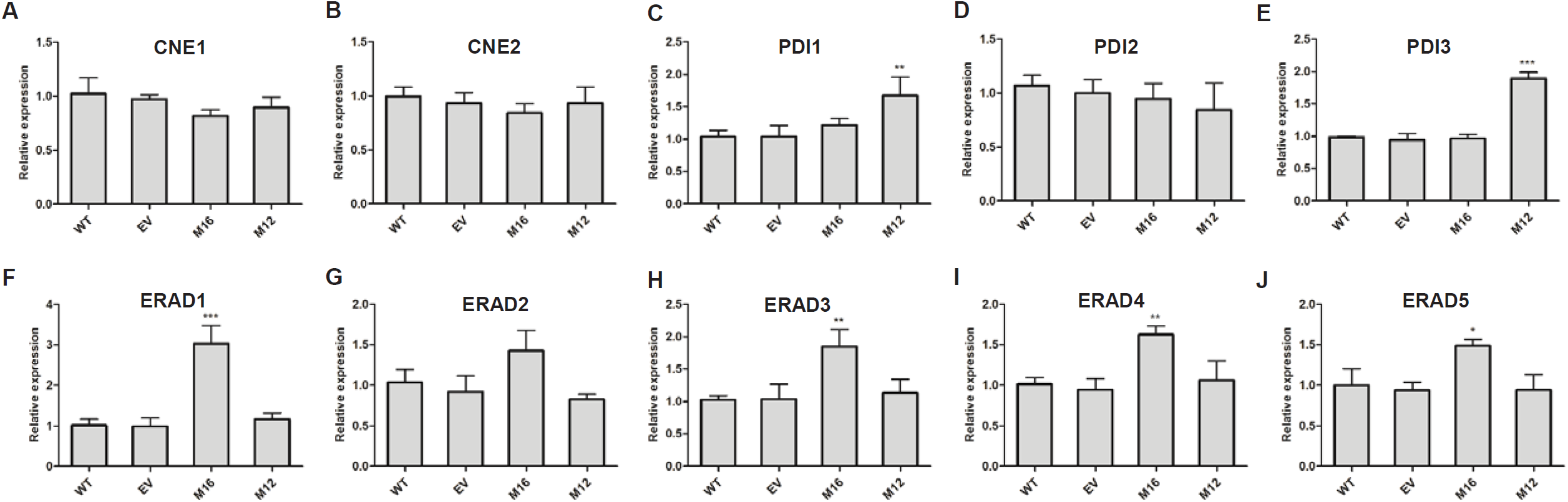
Transcript levels of five unfolded protein response (UPR) and five ER-associated degradation (ERAD) target genes of *P. sojae* in GPI16 and HSP70 mutants. The WT parental strain of *P. sojae* was PS6; EV was the *P. sojae* isolate transformed with an empty vector; M16 was the glycosite mutant of GPI16 with a point mutation at Asn 94 (A-E); M12 was the glycosite mutant of HSP70 with a point mutation at Asn 270 (F-J). (A-B) two major ER chaperones and foldase genes in the UPR pathway; (C-E) four protein disulfide isomerase genes in the UPR pathway; (F-J) five genes in the ERAD pathway. Columns and bars indicate means ± standard deviation (SD) from three biological replicates, and three repeated experiments. An asterisk indicates significant differences based on unpaired Student’s *t* test with the *P* values marked (\**P* < 0.05; \*\**P* < 0.01, \*\*\**P* < 0.001).

## Discussion

Currently, N-glycoproteome data is very limited in phytopathogenic fungi and oomycetes, with the only fungal N-glycoproteome, that of *F. graminearum*, reported in 2016 (13). In the present study, we identified 496 N-glycosylation sites and 355 glycosylated proteins in three developmental stages of *P. sojae*. Similarly, 425 and 516 high-confidence sites were previously mapped in fission and budding yeast (8), while 406 proteins containing 774 sites have been identified in *F. graminearum* (13). According to these results, the proportion of N-glycosylated proteins among the total proteins appears lower in microorganisms than the proportion reported in mammals, using the same cut-off scores for identifying PTM peptides (38). Overall, this study extends the number of known sites and target motifs, and advances the current knowledge surrounding the oomycete N-glycoproteome, providing insight for further exploration of the potential physiological functions of these glycosylated proteins in reproduction and pathogenicity.

Most of the 496 N-glycosylation sites matched with the consensus motif for N-glycosylation N_glyco_-X-S/T (X≠P) as shown in Fig. 2A. This finding is in concordance with the fact that the underlying core “N-glycosylation machinery” consists of few proteins, which are highly conserved in eukaryotic species (11, 51, 52). However, the motif patterns, such as N_glyco_-A, N_glyco_-N, and N_glyco_-D in *P. sojae* (Fig. 2A) differed from glycosylation sites in other species (38, 53). The N_glyco_-N motif has only been previously reported in *F. graminearum* (13), suggesting that these sites may not be conserved among higher organisms. The N_glyco_-G motif has been reported in several species including mammals (8), plants (5), and fungi (5, 13), and the N_glyco_-X-C motif was found in mammals (5, 8). However, these two motifs are absent in oomycetes, indicating an evolutionary divergence in the specificity of N-glycosylation. Overall, these results suggest there are more than one conserved glycosylation motifs in organisms.

N-glycosylation adheres to very rigid topological and sequence constraints, and N-glycosylated proteins have been reported to be mainly located in the cell exterior or in intracellular compartments such as the ER and Golgi (5, 38, 53). The results of this study were consistent with previously reported properties and distributions of N-glycoproteins, with most glycoproteins located in the extracellular matrix or plasma membrane (Fig. 3A), thus supporting the confidence of our N-glycoprotein identifications. However, in contrast to previous studies, we found that the second most common subcellular compartment for glycoproteins was mitochondria in oomycetes (Fig. 3A), especially in the glycoproteins carrying N_glyco_-A, N_glyco_-N, N_glyco_-D, and N_glyco_-S motifs (Fig. 3C), which has only been reported in *F. graminearum* (13), thus reflecting a novel and potentially unique role of N-glycosylation in phytopathogenic microorganisms. In addition, some glycoproteins were localized to the cytoplasm based on curated annotation data (Fig. 3A-3C). An alternative type of N-glycosylation has been identified in bacteria that takes place in the cytoplasm (54-57). This novel cytosolic N-glycosylation is performed by an enzyme structurally different from OST, but which displays the same acceptor site specificity for N_glyco_-X-S/T (11). In light of this finding, we speculated that more kinds of OSTs could be identified in oomycetes due to the unique subcellular localization of glycoproteins.

GO and KEGG pathway analyses revealed that N-glycosylated proteins expressed during the stages examined in this work were involved in ‘carbon metabolism’, ‘TCA cycle’, and ‘fatty acid metabolism’ among others, confirming *P. sojae* glycoproteins belonged to a wide range of biological processes. The calcium ion binding and cation binding functional categories were enriched for the most highly up-regulated N-glycoproteins in the Cy vs My comparison group; this finding indicates that ion flux and homeostasis are the most quantitatively distinct processes between cysts and the mycelia. Calcium is necessary for signaling and function in eukaryotic flagella, and silencing of *Pibzp1*, a regulator of calcium signaling, adversely affected the cyst germination rate and the ability to infect plants in other *Phytophthora* species (58, 59). Notably, KEGG pathway analysis of the differentially expressed N-glycoproteins in the Oo vs My and Oo vs Cy comparisons found that ‘protein processing in endoplasmic reticulum’, ‘other glycan degradation’, and ‘sphingolipid metabolism’ pathways were all enriched for glycoproteins in the oospore stage. N-glycosylation is conducted in the endoplasmic reticulum (10, 11); this result is consistent with the differential increase in expression of glycosylated proteins in oospores compared to the other two stages, indicating a necessary demand for glycoproteins during sexual reproduction.

Deglycosylation analyses and site-directed mutagenesis of the identified glycosites were performed in the current work to validate the results of our N-glycoproteome annotation, and to further interrogate the significant differentially expressed glycoproteins for their role in development and reproduction of *P. sojae*. This study demonstrated that GPI16 and HSP70 are both N-glycosylated at Asn residues in conserved motifs in *P. sojae*, further validating the N-glycoproteome data and confirming the presence and functionality of the N_glyco_-N motif in oomycetes. We also found that both the N_glyco_-X-S/T and the N_glyco_-N motifs are conserved at the genus level, suggesting that these OST recognition sequences have potential for development as fungicide target sites for the control of *Phytophthora*.

The glycosites of these two proteins were mutagenized using CRISPR/Cas9 to ablate their capacity for post-translational glycosylation. Among these, the GPI16 protein participates in GPI anchoring, a ubiquitous mode of post-translational modification in eukaryotic organisms (60). Attachment of GPI anchors to proteins is mediated by the GPI transamidase enzyme complex (60-64). N-glycosylation was necessary for the expression of functional GPI16, as indicated by the decreases in the rate of cyst germination and pathogenicity among glycosite mutants (Fig. 5). The GPI transamidase subunits GPI8, GPI16, and GAA1 form three-component core sub-complexes in *Saccharomyces cerevisiae*, which is conserved in many species (65). Structural modeling suggests that GPI16 may present the substrate protein to GPI8p, and deletion of *GPI16* gene in *Plasmodium berghei* reduces GPIs and GPI-anchored proteins in parasite membranes (66). Further work is needed to determine whether N-glycosylation of GPI16 affects the biosynthesis of GPIs in *Phytophthora*. The second glycosite-mutagenized protein, HSP70 was shown to contribute to cyst germination, oospore production and pathogenicity (Fig. 5). Hsp70 is, by far, a more evolutionarily conserved protein than GPI16 (67, 68), which is consistent with the similarity analysis in this study showing conservation of the N-glycosylation motif between *Phytophtora* and *Pythium* species.

Many lines of evidence have shown that glycoproteins are more stable than their corresponding unglycosylated counterparts, regardless of whether or not the glycosylation is accompanied by major structural changes (69). One proposed explanation is that glycosylation reduces disorder in unfolded proteins, thus increasing the reversibility of the unfolding process and resulting in overall entropic effects that reinforce protein stability. Alternatively, N-linked glycans may be necessary to stabilize some secondary structures that are necessary for protein function or that maintain folded protein conformation (70, 71). To further explore the downstream effects of N-glycosylation on gene expression during different reproductive stages in *Phytophthora*, we measured the relative expression levels of 14 *UPR* and *ERAD* target genes in the glycosylation knockout mutant strains. The mRNA levels of UPR pathway genes *PDI1* and were up-regulated in the HSP70 mutants, while the five *ERAD* genes were all highly expressed in the GPI16 mutants (Fig. 6). Similarly, previous work revealed that in *Trichoderma reesei*, abolishing N-glycosylation of CBH1 led to the induction *UPR* target gene expression, with 5-, 4-, and 2-fold up-regulation of *CNE1, BIP1*, and *PDI1*, respectively (72). These results thus demonstrated that elimination of glycosylation sites can lead to a range of different effects on the folding and stabilization of glycoproteins in oomycetes, though further exploration is needed to characterize the full extent and underlying mechanisms of these effects.

The developmental stages of plant pathogens have been shown to characteristically differ at the transcriptional, post-transcriptional, translational, and post-translational levels (31, 73-75). In particular, N-glycosylation is a post-translational modification critical for fundamental processes underlying growth and reproduction (28-30). In this study, we used PTM-omic approaches to survey changes in N-glycosylation events in the mycelia, cysts, and oospores of *Phytophthora sojae*. Our results indicate that N-glycosylation of proteins occurs extensively throughout the life stages of *Phytophthora* examined in this study. This study is the first, to our knowledge, to confirm the presence of multiple N-glycosylation motifs in oomycetes. This work also demonstrated the crucial role of N-glycosylation in *P. sojae*, and will ultimately provide a large-scale proteomic and proteome-wide glycosylation dataset for *Phytophthora* oomycetes which can be progressively interrogated for insights into the function of glycoproteins in specific processes as well as in macroscopic developmental phenotypes in oomycetes. An increasingly comprehensive understanding of the physiological functions of PTMs in *P. sojae* will aid in the improvement of management and prevention strategies for *Phytophthora*-associated diseases, and may be relevant to homologous PTM-mediated developmental processes in higher organisms.

## Acknowledgments

This work was funded by the National Natural Science Foundation of China (31672052, 31801761) and China Postdoctoral Science Foundation (2019T120160). The authors acknowledge Jingjie PTM BioLabs, Inc. for technical assistance with peptide and protein identification.

## Data availability

Mass spectrometry data have been deposited to the ProteomeXchange Consortium via PRIDE with the data set identifier PXD015294.

## Declaration of Interest Statement

The authors have declared that no competing interest exist.

